# Multiscale entropy is related to iron status in resting state EEG data

**DOI:** 10.64898/2026.08.11.744270

**Authors:** Sarah F. Newbolds, Michael J. Wenger

## Abstract

Dietary iron deficiency in the absence of anemia (IDNA) affects numerous people worldwide, with a wide range of negative effects on brain functioning and cognition. Although studies employing electroencephalography (EEG) have revealed a number of negative effects of IDNA in both the time- and frequency domains, to date there have been no attempts to characterize the effects of IDNA on the temporal dynamics of whole brain interactions. To address this issue, we applied multiscale entropy (MSE) analysis to resting-state EEG data collected from IDNA (n = 21) and iron sufficient (IS, n = 21) women. The MSE analysis on this data revealed that entropy was higher overall for the IS than the IDNA group, with significant differences appearing primarily at longer time scales and under right frontal and left and right parietal electrodes. These results suggest that IDNA may negatively affect long-distance interactions among brain regions and that this could conceivably be a source of diminished cognitive function and neural resilience in IDNA.

## Introduction

Iron deficiency anemia (IDA) affects an estimated 1.2 billion people worldwide, and iron deficiency (ID) in the absence of anemia affects at least twice as many individuals, making ID the most prevalent nutrient deficiency worldwide (Al-Naseem, Sallam, Choudhury, & Thachil, 2021). ID, while a common cause of anemia, is not the only potential cause, as it is possible to be ID without being anemic and to be anemic without having ID. ID is only classified as IDA when hemoglobin levels reach certain values, as outlined by the World Health Organization (hemoglobin [Hb] < 13 g/dL in males, < 12 g/dL in females, < 11 g/dL females during pregnancy Camaschella, 2023), and recent work suggests that the prevalence of ID and IDA may be higher than estimated using these criteria (Mei et al., 2021). Common causes of ID include loss of blood due to menstruation or gastrointestinal problems, pregnancy, low dietary iron intake, and issues with iron absorption (Auerbach, DeLoughery, & Tirnauer, 2025).

ID in the absence of anemia is prevalent in females of reproductive age, affecting an estimated 38% of non-pregnant women in high-income countries (Auerbach et al., 2025). In the US, an analysis of National Health and Nutrition Examination Survey (NHANES) data from 2003 to 2020 estimated that ID affected almost 40% of women aged 12-21 (Weyand et al., 2023). However, this result may be underestimating the true prevalence of ID, as recent research suggests that the NHANES fails to document more than 70% of ID cases due to relying on hemoglobin levels for ID screening (Jefferds et al., 2022; Mei et al., 2021). Based on our experience, more than 40% of college-aged women in the US are ID, anemic, or both, and unaware of their conditions (Wenger, DellaValle, Murray-Kolb, & Haas, 2019). Estimates of ID vary based on population, with a recent study from Lebanon reporting ID in 57.7% of women aged 18-50 (Abuaisha, Itani, El Masri, & Antoun, 2020). There is evidence that ID is under-treated once diagnosed; a statewide study from Minnesota found that 58% of those with ID had not achieved resolution of the condition within three years of diagnosis, and this risk increased among both female and Black populations (Cogan, Meyer, Jiang, & Sholzberg, 2024).

The proper diagnosis and treatment of ID is of critical importance, as it is associated with an increase in all-cause mortality (Schrage et al., 2020), symptoms of depression (Ciulei, Ahluwalia, McCormick, Teti, & Murray-Kolb, 2023), and negative effects on physical and exercise performance (Haas & Brownlie, 2001; DellaValle & Haas, 2011), academics (Scott, De Souza, Koehler, & Murray-Kolb, 2017; More, Shivkumar, Gangane, & Shende, 2013), and cognition (Scott & Murray-Kolb, 2016; Otero, Pliego-Rivero, Contreras, Ricardo, & Fernández, 2004; Lozoff, Jimenez, & Smith, 2006; Wenger, DellaValle, et al., 2019). Additionally, evidence suggests that ID has long-term effects on cognition, with ID in infancy affecting school performance and cognitive functioning through childhood (Congdon et al., 2012; Kumari et al., 2025; Lozoff, Jimenez, Hagen, Mollen, & Wolf, 2000) into early adulthood (Lukowski et al., 2010).

Fortunately, research has found that the cognitive effects of ID, both with and without anemia, can be reversed with iron repletion (Jáuregui-Lobera, 2014). A study of women aged 18-35 found that both addressing ID and IDA women with iron supplementation for 16 weeks produced a five- to seven-fold improvement on tests of attention, memory, and learning (Murray-Kolb & Beard, 2007). Our work has demonstrated comparable results with dietary sources of iron. In a study of college-aged women in Rwanda (Wenger, Rhoten, et al., 2019), non-anemic women with ID who consumed an iron biofortified bean for 128 days showed shorter reaction times and higher scores on memory and attentional tasks relative to those who consumed a control bean. Additional research has found that women of reproductive age in India with ID scored higher on tasks of perception, attention, and mnemonics after receiving supplementation with double-fortified salt (Wenger et al., 2017), and IDA and ID adolescents who consumed iron biofortified grain showed improved cognitive performance (Wenger, Murray Kolb, Scott, Boy, & Haas, 2022) relative to those who consumed a non-fortified grain. Even in a group of adolescent girls with a relatively low prevalence of ID, consumption of iron fortified lentils (Barnett, Wenger, Yunus, Jalal, & DellaValle, 2023) improved overall cognitive performance.

### Iron, the brain, and brain dynamics

Iron plays several roles in the central nervous system, including the myelination of neurons, energy metabolism, neuro- and synaptogenesis, oxygen transport and storage, and neurotransmitter synthesis and regulation (Beard & Connor, 2003; Gao et al., 2025). Work in animal models has demonstrated significant effects of ID on DA functioning, including reductions of dopamine D1 and D2 receptors and dopamine transporters (DAT) in the caudate putamen and nucleus accumbens, leading to significant decreases in central DA neurotransmission (Erikson, Jones, & Beard, 2000; Erikson, Jones, Hess, Zhang, & Beard, 2001; Pino et al., 2017; Unger, Wiesinger, Hao, & Beard, 2009; Youdim, Ben-Shachar, & Yehuda, 1989). ID is also responsible for higher levels of extracellular DA in the striatum, providing evidence that ID negatively affects DA reuptake (Erikson et al., 2000; Nelson, Erikson, Piñero, & Beard, 1997). The effects of ID on DA functioning has not been widely studied in humans, as direct measures of DA are invasive. However, a recent study by our lab documented a reliable relationship between retinal activity activity (an indirect meaasure of DA), and provided evidence that DA mediates the association between iron levels and measures of cognition in college-aged women (Newbolds & Wenger, 2024).

Research has also demonstrated that ID, even in the absence of anemia, causes significant changes in brain activity as measured by electroencephalography (EEG). In fact, some of the earliest work on ID and IDA in humans documented the effects on EEG (Tucker & Sandstead, 1981; Tucker, Sandstead, Penland, Dawson, & Milne, 1984). Otero et al. (1999) found that children aged 6-12 with ID showed slower activity in the EEG power spectrum than iron sufficient (IS) children. Additional research on ID children demonstrated that iron supplementation can normalize P300 event-related potential (ERP) responses (2004), and return diminished ERP amplitudes to normal, as well as improve working memory (Otero, Pliego-Rivero,

Porcayo-Mercado, & Mendieta-Alcántara, 2008). Our studies have also shown that dietary supplementation with iron can improve EEG outcomes. IDA and ID adolescents in India who consumed iron biofortified grain showed improvements in EEG measures of attention, including *α* and *γ* power, and the amplitude of specific ERPs (Wenger et al., 2022). Similarly, college-aged women with ID who consumed iron biofortified beans showed an increase in the peak amplitude of specific ERPs, with mixed results in the *α* and *γ* frequency bands suggested (Wenger, Rhoten, et al., 2019). A recent study by Rhoten et al. (2025) found that women with ID demonstrated large differences as compared to IS women on EEG measures of brain dynamics and neural efficiency (Wenger, Townsend, & Newbolds, 2025).

Critically, in two separate studies (Newbolds & Wenger, 2024; Rhoten et al., 2025) we have shown that it is possible to reliably distinguish those with from those without ID solely on the basis of brain activity. This research suggests that ID, with and without anemia, has the potential to create distinct brain states, defined as specific and recurring whole-brain neural activity pattern emerging from physiological or cognitive processes (Greene, Horien, Barson, Scheinost, & Constable, 2023). This in turn suggests that it may be possible to identify patterns of brain activity at multiple time scales that distinguish ID from iron sufficient (IS) nervous systems. In this research, we provide evidence that of this as measured by multiscale entropy (MSE).

The brain can be considered as a dynamic, non-linear system, and as a result, complexity measures typically applied to such systems are increasingly useful for studying the complex neural interactions underlying cognitive functioning, including in the context of mental disorders, disease states, and aging (Cofré & Destexhe, 2025; Takahashi, 2013). Multiscale entropy (MSE) is an especially effective approach for quantifying complex brain activity on multiple time scales (Costa, Goldberger, & Peng, 2002). MSE has been used to study aging and neurodegenerative disease, where healthy individuals tend to shower higher levels of complexity, and thus, entropy, in neural data than their affected counterparts (Lau, Pham, Chen, & Makowski, 2022; Zúñiga et al., 2024). Here, higher levels of complexity, thus entropy, can be interpreted in terms of the number of possible states that neural systems can occupy, with more states—higher complexity—indicating greater potential for resilience. In a group of older adults, Iinuma et al. (2022) found that those in a high cognitive functioning group demonstrated higher levels of MSE at frontal, parietal, and temporal regions than individuals in a lower cognitive functioning group. A study by Fan et al. (2018) demonstrated that decreasing MSE with associated with the severity of AD. Despite being performed on only 10 s of resting state EEG data, the MSE analysis accurately distinguished between individuals with severe AD and normal controls. The present study sought to determine whether ID, in the absence of anemia, is associated with lower complexity in otherwise healthy college-aged women, as measured using MSE analysis. This would suggest that ID results in neural systems with lowered levels of resilience, underscoring the systemic effects of ID.

## Methods

The project used unpublished data from an existing dataset collected in a previous study (Newbolds & Wenger, 2024). Full methodological details can be found in that paper and are repeated here in abbreviated form.

### Participants

Participants in this study were 42 female students at the University of Oklahoma, aged 19-29. The sample was 71% White and 29% Asian. Participants were identified as iron deficient non-anemic (IDNA, *n* = 21) if their serum ferritin (sFt) < 12 ng/mL and their hemoglobin (Hb) > 12 g/dL. Participants were identified as iron sufficient (IS, *n* = 21) if their sFt > 20 ng/mL and their Hb > 12 g/dL. Participants were matched on age and ethnicity. Women whose Hb < 12 g/dL were identified as IDA and excluded from participation.

### Task and concurrent EEG

Participants completed a single session in a dimly lit sound-attenuated and electrically shielded chamber. Participants were seated in an adjustable height chair and placed their heads in a chinrest located 72 cm from the computer display. EEG data were collected using high-density (128 channel) electrode nets (Magstim/EGI, Eugene, OR). Data were acquired using a 128-channel Net Amps 300 amplifier and were digitized at a sampling rate of 1 kHz. Impedances were kept at or below 50 KΩ during testing. Data were collected with online filters set at 0.1 Hz (highpass), 70 Hz (lowpass), and 60 Hz (notch). In addition, electrodes (RetEval, (LKC Technologies, Gaithersburg, MD) for collecting the electroretinogram (ERG) were applied immediately below each eye and data were acquired using the amplifier used for the EEG. The EEG data analyzed here were acquired while participants sat for five minutes in an alert resting state. During the recording, participants were asked to focus on a small character on the center of the computer screen that changed orientation every 15 sec. EEG data preprocessing was performed using EEGLab (Delorme & Mackeig, 2004; Delorme, Jung, Sejnowski, Makeig, et al., 2005) for Matlab (Mathworks, Natick MA) following a standard preprocessing pipeline (Delorme, 2023).

### MSE analysis

Prior to analysis, a total of 18 electrodes located along the lowest run of electrodes on the 128 channel net (those approximately at the hair line) were removed from the data set for each participant, as these electrodes typically show the lowest signal-to-noise ratio. A total of five 1 sec segments of data from the resting period for each participant were used in the analyses. Data were downsampled to 500 Hz prior to MSE analysis.

MSE was calculated for each participant following Costa et al. (2002):

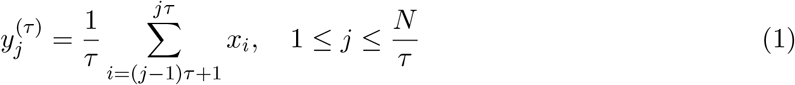

where 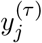 is the *j*th point in the coarse-grained time series at scale factor *τ, j* indexes the consecutive non-overlapping windows, *τ* is the scale factor (i.e., the number of consecutive samples averaged to produce each coarse-grained data point), *x*_*i*_ is the *i*th sample in the original EEG time series, and *N* is the total number of samples in the original time series. Estimating the complexity in this way allows for a more comprehensive analysis of brain-wide dynamics across multiple time scales and further provides insight into the complexity of the signal over distances (Costa et al., 2002; Iinuma et al., 2022). The data for each participant were averaged across the five segments at each time scale, and the time scale ranged from 4 ms (*τ* = 1) to 80 ms (*tau* = 20).

## Results

The full set of behavioral and electrophysiological results are available in Newbolds and Wenger (2024). In brief, cognitive performance and aspects of the ERG tracked iron status, with aspects of the ERG mediating the relationship between iron status and cognitive performance and suggesting that the ERG may function as a reliable proxy for DA status in the context of ID.

To test for group differences (IDNA vs. IS), we applied a *t*-test with a family-wise false discovery rate (Benjamini & Hochberg, 1995) to the MSE scores. The results showed that the entropy was higher overall for the IS than the IDNA group (Figure 1(a)). Significant differences appeared at longer time scales, with *τ* = 14 *−* 20, corresponding to 56-80 ms time intervals (Figure 1(b)). Significant differences were also found under clusters of electrodes above the right frontal lobe (electrodes 5-10), left parietal lobe (electrodes 31-66), and right parietal lobe (electrodes 96-122). The time range involved as well as the spatial separation of the clusters suggests effects associated with long range projections, such as those that may be involved in various distributed attentional networks (e.g., Menon & D’Esposito, 2022).

**Figure 1.**
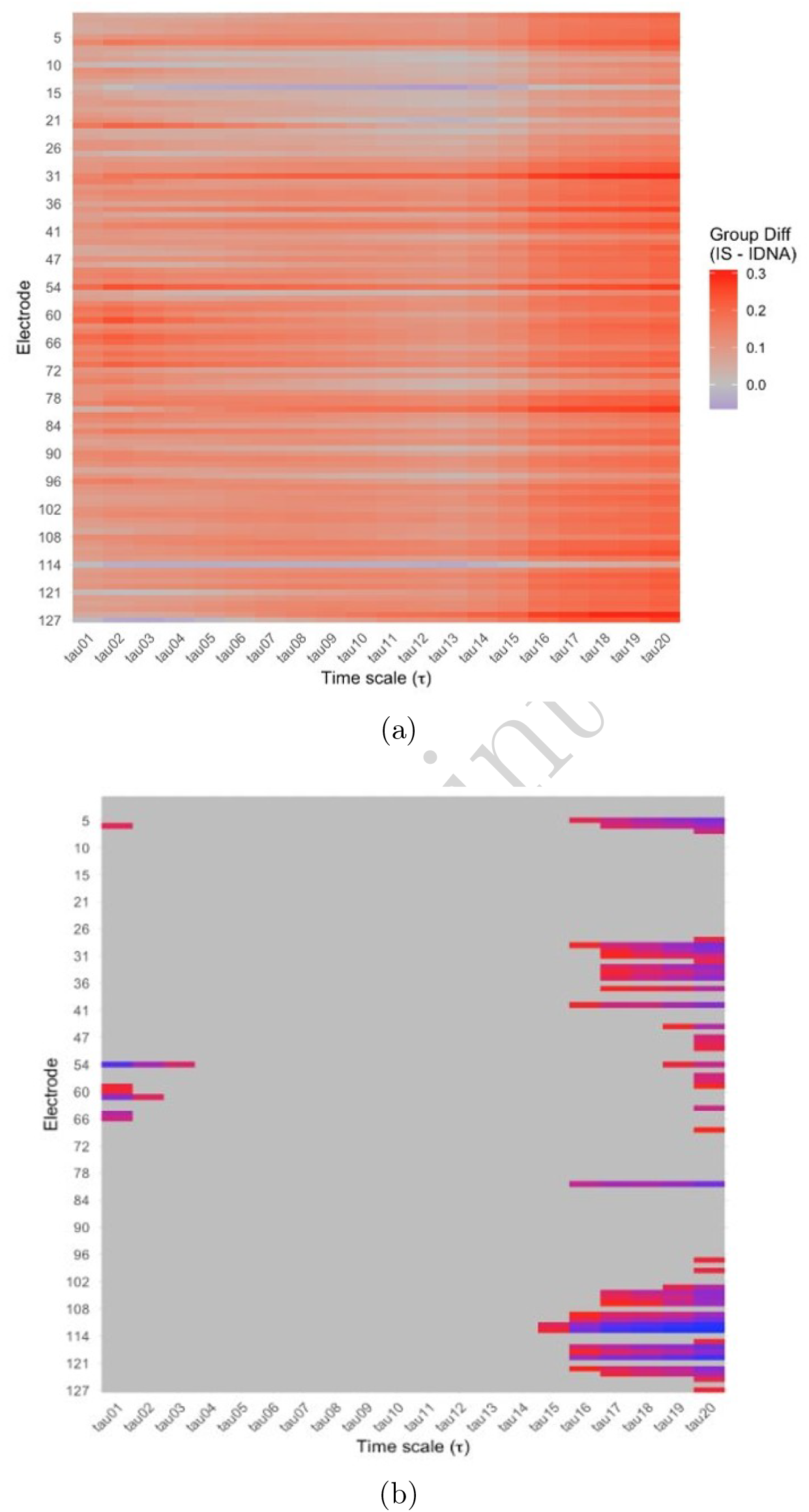
(a) Group differences in MSE, with red indicating IS > IDNA; (b) group differences thresholded at *p* ≤ 0.001, with colder colors indicating smaller *p*-values. MSE was higher overall for the IS than the IDNA group, with significant differences appearing at longer timescales and under right frontal and right and left parietal electrodes.

## Discussion

This study applied an analysis of multiscale entropy to resting-state EEG data to test for differences in neural dynamics in IS versus IDNA women. The results of the analysis demonstrated a significant difference in complexity between the two groups, with higher complexity found in the IS group. Significant differences were found primarily at longer time scales, and under electrodes corresponding to the right frontal and right and left parietal lobes. This suggests that ID is associated with a reduction in neural complexity and, by extension, neural resilience, and that this reduction involves broadly distributed regions of the brain, consistent with accumulating evidence (e.g., Otero et al., 2004; Tucker et al., 1984; Wenger, Rhoten, et al., 2019; Rhoten et al., 2025) of the broad impact of ID without anemia on neural function.

The results of this analysis were similar to those reported by Iinuma et al. (2022), who performed an MSE analysis on individuals aged 65-85 in two groups defined by level of cognitive functioning. That study found that individuals in the high cognitive functioning group had significantly higher MSE than those in the low cognitive functioning group. Additionally, these complexity differences were found in slower time scale regions, with significant differences in electrodes over the frontal, parietal, and temporal lobes of the brain. Iinuma et al. (2022) suggested that slower time scales reflect long-distance neural interactions among different brain regions that are needed to achieve higher cognitive functioning. The results of the current work suggest that IDNA similarly may negatively affect long-distance interactions among brain regions and that this could conceivably be a source of diminished cognitive function in IDNA.

A potential mechanism for the observed reduction of complexity in the IDNA women could be the effect of iron deficiency on neurotransmitter synthesis and regulation, including dopaminergic communciation. Previous research has suggested that ID, even in the absence of anemia, can disrupt normal dopaminergic functioning in the brain (e.g., Beard & Connor, 2003; Newbolds & Wenger, 2024; Unger et al., 2009). Dopamine plays a central role in the frontalparietal control network (FPCN), which is heavily implicated in a variety of aspects of cognition (Dang, O’Neil, & Jagust, 2012; Marek & Dosenbach, 2018; Menon & D’Esposito, 2022).

Overall, the results of this study documented a significant difference in complexity, as measured by MSE, in IDNA versus IS women. The results suggest that IDNA may negatively affect long-distance connections among distributed brain regions, leading to diminished cognitive performance and overall neural resilience. These results are the first application of complexity measures to study the effects of IDNA on brain function in otherwise healthy women and support the hypothesis that IDNA negatively impacts whole-brain dynamics.

## Declarations

### Ethics approval

This research was reviewed and approved by the Institutional Review Board of The University of Oklahoma. All work was performed in accordance with the ethical standards as laid down in the 1964 Declaration of Helsinki and its later amendments.

### Consent to participate

Written informed consent to participate was obtained from all participants at the outset of their participation.

### Consent to publish

Written informed consent to publish de-identified data was obtained from all participants at the outset of their participation.

### Availability of data and materials

All data and experimental materials are available upon reasonable request to the corresponding author.

### Code availability

All code developed for running the studies and analyzing the data are available upon reasonable request from the corresponding author.

### Authors’ contributions

SFN: data analysis, writing and revising the manuscript. MJW: conception, design, writing and revising the manuscript. MJW: funding.

